# Lack of paternal silencing and ecotype-specific expression in head and body lice hybrids

**DOI:** 10.1101/2023.05.07.539726

**Authors:** Hollie Marshall, Andrés G. de la Filia, Ross Cavalieri, Eamonn B. Mallon, John M. Clark, Laura Ross

## Abstract

Paternal genome elimination (PGE) is a non-Mendelian inheritance system in which males develop from fertilised eggs but their paternally-inherited chromosomes are eliminated before or during spermatogenesis. Therefore, PGE males only transmit their maternally inherited set of chromosomes to their offspring. PGE has been described in numerous arthropod species, many of which are pests or parasites, posing a severe economic burden on crop production and/or with implications for human health. In order to understand how PGE has evolved on the molecular level, to potentially develop novel control strategies, we need to examine species which display basal forms of PGE. The human louse, Pediculus humanus, represents an ideal model system to understand the molecular underpinnings of PGE. In this study we analysed parent-of-origin allele specific expression patterns in male offspring of crosses between head and body lice ecotypes. We have shown that hybrid adult males of P. humanus display biparental gene expression, which constitutes the first known case of a species with PGE in which genetic activity of paternal chromosomes in the soma is not affected by embryonic heterochromatinization or (partial or complete) elimination. We have also identified maternally-biased genes (potentially imprinted genes) which may be involved in the elimination of paternal chromosomes during spermatogenesis. Finally, we have identified genes which show ecotype-specific expression bias. Given the low genetic diversity between ecotypes this is suggestive for a role of epigenetic processes in ecotype differences. These findings have implications for models of pediculicide resistance in human lice and for the development of novel epigenetic-mediated control strategies.

## Introduction

Many organisms follow Mendelian inheritance where offspring receive a random chromosome set from each parent. However, Mendel’s Laws are broken by 1000s of species where the inheritance of the maternal or paternal chromosomes is biased. Paternal genome elimination (PGE) is one such system. It is widely distributed across arthropods, in which males develop from fertilised eggs but their paternally-inherited chromosomes are eliminated before or during spermatogenesis. Therefore, PGE males only transmit their maternally inherited set of chromosomes to their offspring (Normark, 2003; Burt and Trivers, 2006; Gardner and Ross, 2014). In PGE species where elimination of paternal chromosomes takes place early in development (embryonic PGE), such as mesostigmatid mites (Nelson-Rees *et al*., 1980; Norton *et al*., 1993) or diaspidid scale insects (Nur, 1980; Ross *et al*., 2010), males become true haploids and therefore only express maternal alleles. In some PGE species where elimination is delayed until spermatogenesis (germline PGE), such as Scolytinae beetles (Brun *et al*., 1995), neococcid scale insects (Brown and Nur, 1964; Bongiorni and Prantera, 2003; Ross *et al*., 2010) or booklice (Hodson *et al*., 2017), paternal chromosomes are retained throughout development, but become tightly condensed during embryogenesis. As a result, paternal chromosomes remain transcriptionally inactive (Berlowitz *et al*., 1968) and expression of maternal alleles usually predominates. One of the most intriguing aspects of the evolutionary history of PGE is the repeated convergence towards these adaptations to reduce or completely prevent expression of paternal chromosomes in males in the different groups where PGE has independently emerged.

In order to understand how PGE has evolved on the molecular level, we need to examine species which display potentially basal forms. The human louse *Pediculus humanus* is one such species (McMeniman and Barker, 2006; de la Filia *et al*., 2018). Similar to other PGE taxa, males have a strongly modified spermatogenesis: the first meiotic division is followed by a series of mitosis to form a 32/64 cell cyst, of which only half (those containing maternal chromosomes) continue to develop as active spermatozoa, while the other degenerate *in situ* (Hindle and Pontecorvo, 1942; Bressa *et al*., 2015). However, none of the somatic phenomena associated to PGE—heterochromatization, elimination of single paternal chromosomes or the whole paternal set during early development—have ever been described in human lice, despite extensive cytogenetic research. It therefore remains unknown as to whether the paternal chromosomes in males are transcriptionally active or not. From an evolutionary perspective, human lice could then represent the most basal form of PGE under the evolutionary arms race hypothesis, prior to the emergence of any putative maternal adaptations to counteract resistance of paternal alleles to germline elimination (Herrick and Seger, 1999; Ross *et al*., 2010), consistent with the high rate of paternal allele leakage found in de la Filia *et al*. (2018). If this is the case, *P. humanus* would be a promising system to directly search for genes or chromosomal regions with parent-of-origin-specific expression which could be directly involved in the induction—or deterrence—of paternal chromosome elimination.

Moreover, somatic patterns of gene expression in *P. humanus* are also interesting beyond fundamental theories on intragenomic conflict and evolution of non-canonical genetic systems. Many PGE species with paternal chromosome silencing are pests or parasites that pose a severe economic burden on crop production (Gill and Kosztarab, 1997; Damon, 2000). *P. humanus* is a widespread human ectoparasite with serious consequences (Clark *et al*., 2013). Head and body lice are two distinct ecotypes of *P. humanus*, differing in morphological and behavioural traits driven by ecological factors (Light *et al*., 2008; Veracx and Raoult, 2012). Body lice constitute a serious health threat as vectors of severely pathogenic bacteria (Roux and Raoult, 1999) and head lice infestations have been estimated to cause costs of hundreds of million dollars every year (Hansen and O’Haver, 2004). Recently, increasing resistance to available treatments has become a major challenge to human louse control (Burgess, 2004; Durand *et al*., 2012; Clark *et al*., 2015; Clark, 2018), creating the need for novel resistance-proof management programmes. The overlooked transmission patterns of PGE in both ecotypes could inform novel pediculicide-free management strategies. Yet, a full characterization of the form of PGE present in *P. humanus* is necessary to determine whether existing models of resistant evolution in haplodiploid taxa (Crozier, 1985; Caprio and Hoy, 1995; Denholm *et al*., 1998; Carrière, 2003) can be applied.

In this study we analysed parent-of-origin allele specific expression patterns in F1 male offspring of crosses between head and body lice. The transcriptomic profiles of both ecotypes are highly similar, with low levels of inter-ecotype divergence in nucleotide sequences and gene expression levels (Olds *et al*., 2012). Therefore, individuals from both ecotypes can be easily crossed in laboratory conditions, yielding viable and fertile offspring (Busvine, 1978). Using the parent-of-origin data, we ask 1) is there expression of paternal chromosomes in hybrid males? 2) is there ecotype-specific expression in hybrid males? and 3) does ecotype-specific expression in hybrid males match parental gene expression patterns for each ecotype?

## Methods

### Experimental populations and inter-ecotype crosses

A series of intraspecific crosses were set up using individuals from the head louse strain SF-HL and the body louse strain Frisco-BL. The SF-HL colony was established in 2002 from head lice collected from ∼20 infested children in Plantation, Miami, and Homestead (FL, U.S.A.). Approximately 50 males and 50 females were used to temporarily establish a colony on human volunteers (Takano-Lee et al. 2003). Fertile eggs from Homestead were added to the colony at least three times between 2002 and 2006. Approximately 30–50 eggs were introduced each time. The colony was placed on an in vitro rearing system in 2006 (Yoon *et al*., 2006). The Frisco-BL colony of human body lice was originally collected from nine homeless individuals in San Francisco (CA, U.S.A.) by Dr. Jane Koehler (University of California San Francisco Medical Center, San Francisco, CA, U.S.A.) in December 2008. Both colonies have been maintained by the Clark Laboratory at the University of Massachusetts-Amherst on human blood using the same in vitro rearing system (Yoon *et al*., 2006) under environmental conditions of 30 °C, 70% relative humidity and an LD 16:8 h photoperiod in rearing chambers (University of Massachusetts-Amherst Institutional Review Board approval no. E1404/001-002).

Parental generations (F0) were established by isolating 6 sexually immature third instar males and females from both colonies. F0 males from each ecotype were kept in common cages until sexual maturity, while F0 females were transferred to individual cages. Since sexually mature louse females can lay unfertilised eggs that do not hatch (Bacot, 1917), matings were delayed until females laid a first batch of eggs to confirm virginity. After 1-2 days, these eggs were removed from female cages to be incubated for a minimum of 10 days and then an F0 male from the other ecotype was introduced. Females were allowed to lay eggs for 15 days before removal of the mating pair. In total, five head louse (HH) female x body louse (BB) male crosses (HB1, HB2, HB3, HB5, HB6) and four BB female x HH male crosses (BH1, BH4, BH5, BH6) were successful. Adult F1 males were collected in RNAlater after a 24 h period of starvation. In addition, to generate genomic and transcriptomic data from HH and BB ecotypes, adult individuals were directly isolated from the colonies (only males for RNA-seq, from both sexes for DNA-seq) and collected in RNAlater.

### RNA and DNA extraction and sequencing

RNA was extracted from a single F1 male per cross. Males were removed from RNAlater, washed twice in ice-cold 1X PBS and ground in 400*μ*l of TRIzol (Invitrogen). Total RNA samples were isolated with a PureLink RNA purification kit (Thermo Fisher Scientific, USA), purified with RNA Clean & ConcentratorTM-5 (Zymo Research, USA) and validated using the Bioanalyzer RNA 6000 Nano kit (Agilent). TruSeq stranded mRNA-seq libraries were generated by Edinburgh Genomics (UK) and sequenced on the Illumina NovaSeq platform (S2 flowcell, 50bp paired-end reads).

In addition to F1 RNA-seq, we also generated two replicates of RNA-seq data from pools of 10 adult BB and HH males from the same strains. Both HH and one of the BB samples were sequenced on the Illumina HiSeq 4000 (75 bp paired-end). The second BB sample was sequenced with the F1 RNA-seq. For whole genome re-sequencing of parental strains, DNA was extracted using a DNeasy Blood & Tissue kit (Qiagen, The Netherlands) from pools of 10 adult individuals. TruSeq DNA Nano gel free libraries (350bp insert) and sequencing on the Illumina HiSeq X (150bp paired-end) were performed by Edinburgh Genomics.

### Generation of N-masked genomes

Whole genome re-sequencing data were quality checked using fastqc v.0.11.5 (Andrews, 2010) and aligned to the reference genome (JCVI_LOUSE_1.0, Kirkness *et al*. (2010)) using bowtie2 v.2.3.5.1 (Langmead and Salzberg, 2013) in *–sensitive* mode. Data were deduplicated and read groups added using picard v.2.6.0 (BroadInstitute, 2019). SNPs were called using freebayes v.1.1.0 (Garrison and Marth, 2012) and filtered using vcftools v0.1.14 (Danecek *et al*., 2011), requiring a minimum depth of 10 reads and a minimum quality score of 20. Bedtools v.2.28.0 (Quinlan and Hall, 2010) was then used to identify homozygous alternative SNPs unique to each ecotype, this resulted in 204,411 SNPs unique to HH and 156,114 SNPs unique to BB. An N-masked genome was then created using all of these SNPs via the *maskfasta* command from Bedtools v.2.28.0 (Quinlan and Hall, 2010). The creation of an N-masked genome avoids later mapping bias to the reference allele.

### Differential gene expression between ecotypes

A differential gene expression analysis was carried out between the parental ecotypes of head (HH) and body (BB) lice. RNA-seq data were quality checked with fastqc v.0.11.5 (Andrews, 2010) and trimmed using CutAdapt v.1.11 (Martin, 2011). Reads were aligned to the N-masked genome created above and transcript abundances were calculated using RSEM v.1.3.0 (Li and Dewey, 2011), implementing STAR v.2.7.1 (Dobin *et al*., 2013) with standard parameters. DESeq2 v1.28.1 (Love *et al*., 2014) was used to determine differentially expressed genes between ecotypes. A gene was considered differentially expressed if the corrected p-value was <0.05 (adjusted for multiple testing using the Benjamini-Hochberg procedure (Benjamini and Hochberg, 1995)) and the log2 fold-change was >1.5.

### Identification of parent-of-origin and ecotype-of-origin expression

Male hybrid F1 offspring RNA-Seq data were quality checked using fastqc v.0.11.5 (Andrews, 2010) and trimmed with CutAdapt v.1.11 (Martin, 2011). Reads were aligned to the N-masked genome created above using RSEM v.1.3.0 (Li and Dewey, 2011), implementing STAR v.2.7.1 (Dobin *et al*., 2013) with standard parameters. Bam-readcount v.1.0.1 (https://github.com/genome/bam-readcount) was then used to count the number of reads that map to each of the four nucleotide bases for all positions that contain a homozygous alternative SNP in either head or body lice, identified above from the whole genome re-sequencing data. Count data for the parental nucleotide at each of these positions was then extracted and all positions which fell within a gene were annotated with a gene id using a custom R script. Only genes which had a SNP covered by a minimum of 10 reads in at least two replicates per cross direction were kept for further analysis. Genes showing parent-of-origin or ecotype-of-origin expression were determined using a logistic regression model in R, with a quasibionimal distribution to account for overdispersion. Correction for multiple testing was carried out using the Benjamini-Hochberg method (Benjamini and Hochberg, 1995). Genes were determined as showing parent-of-origin expression if the maternal/paternal allelic ratio (head/body lice allelic ratio for ecotype-of-origin expression) corrected p-value was <0.05 and the parental expression proportion was >0.6 in both cross directions. All custom scripts are available online, see Data Accessibility section.

### Additional analyses

A reciprocal protein blast was carried out to identify the genes involved in DNA methylation maintenance (DNMT1) and establishment (DNMT3a) using 321 insect DNMT1 gene sequences and 110 insect DNMT3a gene sequences from http://v2.insect-genome.com/Pcg, using blastp v2.2.3 (Camacho *et al*., 2009). DNA methylation appears to be present in *P. humanus* based on the CpG observed/expected proxy (Bewick *et al*., 2017). We also then examined this specifically within exons and introns of all genes following custom scripts (https://github.com/MooHoll/cpg_ observed_expected).

Gene ontology (GO) terms were annotated for 4,606 genes out of 9,830 with gene expression data using eggNOG-mapper v.2.0.0 with standard parameters (Cantalapiedra *et al*., 2021). GO enrichment was carried out in R using GOstats v2.56.0 (Falcon and Gentleman, 2007) which implements a hypergeometric test with Benjamini-Hochberg correction for multiple testing (Benjamini and Hochberg, 1995). GO biological processes were classed as over represented if the corrected p-value was <0.05. REVIGO (Supek *et al*., 2011) was then used to visualise GO terms. Drosophila orthologs were determined using VectorBase (Amos *et al*., 2022).

## Results

### Differential gene expression between ecotypes

Using RNA-Seq from the parental ecotypes, we have identified differentially expressed genes between head and body lice males. We find the majority of the variability within the data is between the two body lice replicates, however, 45% is explained by ecotype (Fig.1a). Whilst the majority of genes show similar expression levels between ecotypes (Fig.1b), we do find 47 differentially expressed genes (adjusted p-value <0.05 and log2 fold-change >1.5, supplementary 1.0.0). Significantly more of these are upregulated in body lice compared to head lice (*X-squared* = 9.383, *df* = 1, *p* <0.01, Fig.1c) with two genes being limited to body lice (i.e. with zero expression in head lice, Fig.1d: PHUM135320 - *hypothetical protein* and PHUM619270 - *chloride intracellular channel*).

**Figure 1:**
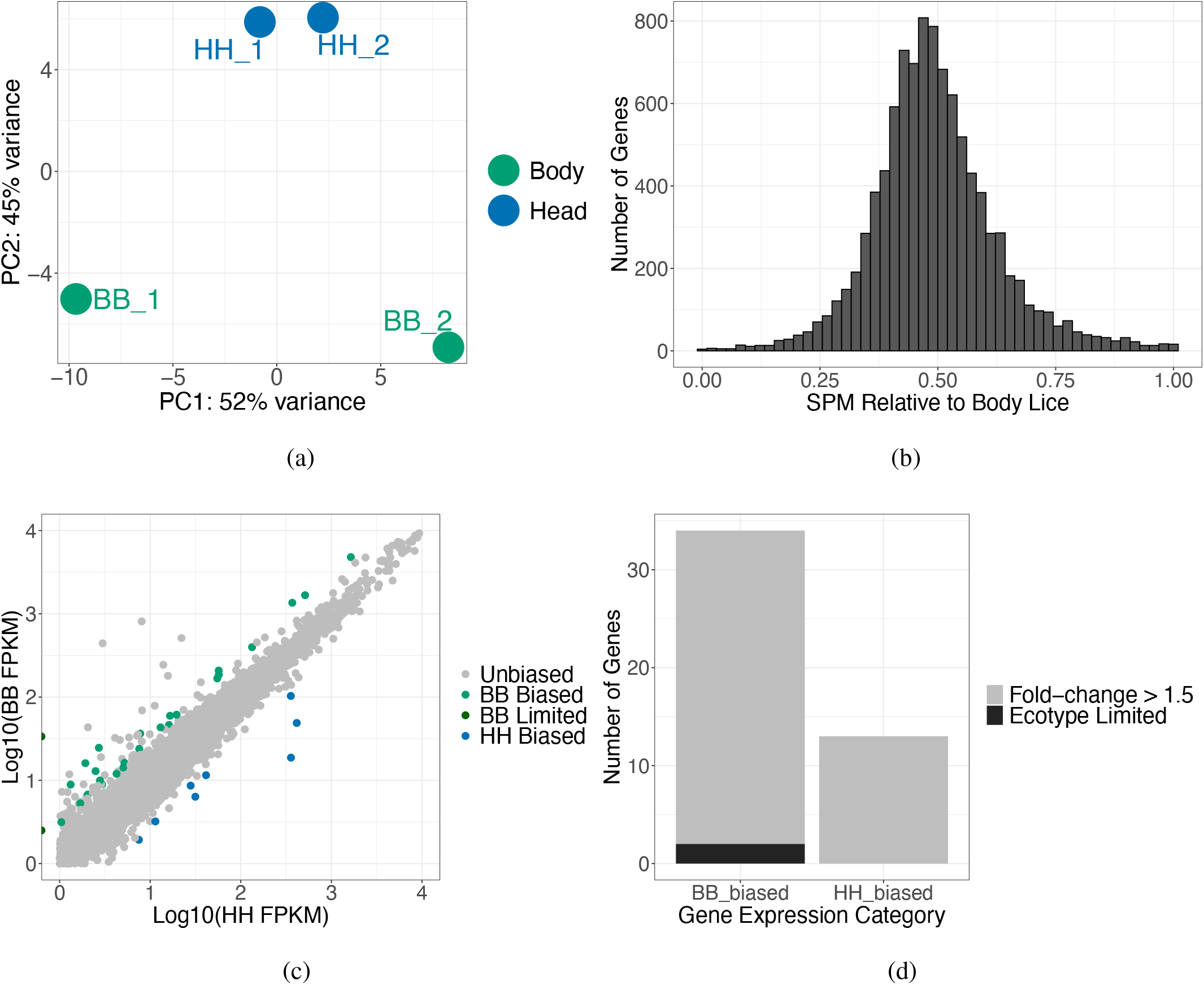
(a) PCA plot based on the expression of all genes in the head and body lice RNA-Seq (n = 9,830). (b) Histogram of the SPM (measure of specificity (Xiao *et al*., 2010), calculated as body lice FPKM squared divided by body lice FPKM squared plus head lice FPKM squared) per gene (n = 9,830) showing a large number of genes are equally expressed between ecotypes. (c) Scatter plot of the log10 fragments per kilobase of transcript per million mapped reads (FPKM) of all genes (n = 9, 830). Significantly differentially expressed genes (corrected p-value <0.05 and log2 fold-change >1.5) are coloured by ecotype and level of differential expression, unbiased genes are shown in grey. (d) Stacked bar plot showing the number of ecotype-biased genes. Ecotype-limited genes referring to those with zero expression in one ecotype.

GO terms enriched for differentially expressed genes between ecotypes are involved in a variety of processes including “*adult feeding behaviour*” (GO:0008343) and “*chitin-based cuticle development*” (GO:0040003). Additionally, multiple GO terms associated with epigenetic processes are enriched, including “*DNA methylation on cytosine within a CG sequence*” (GO:0010424), “*DNA methylation-dependent heterochromatin assembly*” (GO:0006346) and “*histone H3-K27 acetylation*” (GO:0043974), supplementary 1.0.1. Of the two genes with body louse limited expression, one has no GO annotations (PHUM619270), the second (PHUM135320) is involved in wound healing (GO:0042060, GO:0009611) and response to stress (GO:0006950). Two of the most upregulated genes in head lice compared to body lice are involved in nervous system development (PHUM494820) and immune response (PHUM365700 - *Defensin 1*).

In order to further explore the potential involvement of epigenetic processes in mediating ecotype differences, we identified genes involved in DNA methylation maintenance and establishment, DNMT1 and DNMT3a in *P. humanus*. We find similar levels of expression for both genes between ecotypes (Supplementary Fig.S1, DNMT1 (PHUM556380) adjusted p-val = 0.99, DNMT3a (PHUM041250) adjusted p-val = 0.95). Previous research has suggested DNA methylation is present in *P. humanus*, based on the the CpG observed/expected distribution (Bewick *et al*., 2017). We checked this distribution in exons and introns and find DNA methylation is likely enriched in exons in *P. humanus* (Fig.S1), as is seen across arthropods (Lewis *et al*., 2020).

Finally, we also examined differentially alternatively spliced genes between ecotypes using the IsoformSwitchAnalyzeR v.3.16 R package (Vitting-Seerup and Sandelin, 2019). Differences in alternative splicing have previously been found between head and body lice and are suggested to contribute to ecotype phenotypic differences (Tovar-Corona *et al*., 2015). Using the RNA-Seq in our study, we were unable to identify any differentially alternative spliced genes between ecotypes. Whilst we cannot corroborate the previous study, we believe this is due to the lack of annotation of alternative transcripts within the current genome annotation, rather than ecotypes actually lacking differentially alternatively spliced genes.

### Parent-of-origin expression in F1 hybrid offspring

Males of most species with PGE tend to have eliminated or silenced paternal chromosomes within somatic tissue, meaning predominately maternal alleles are expressed. Using reciprocal crosses of head and body lice, we were able to identify parent-of-origin expression in hybrid male offspring of *P. humanus* for 482 genes (supplementary 1.0.2). Surprisingly, we find the majority of these are biparentally expressed, i.e. the paternal allele is not silenced (Fig. 2a). However, there is a skew towards more maternal expression in both crosses generally (except for genes with head lice specific bias, discussed below), Fig. 2b. We also find 18 genes with significant maternal expression bias, six of which show complete silencing of the paternal allele (Fig. 2c). None of these significant maternally biased genes are differentially expressed between pure head and body lice males.

**Figure 2:**
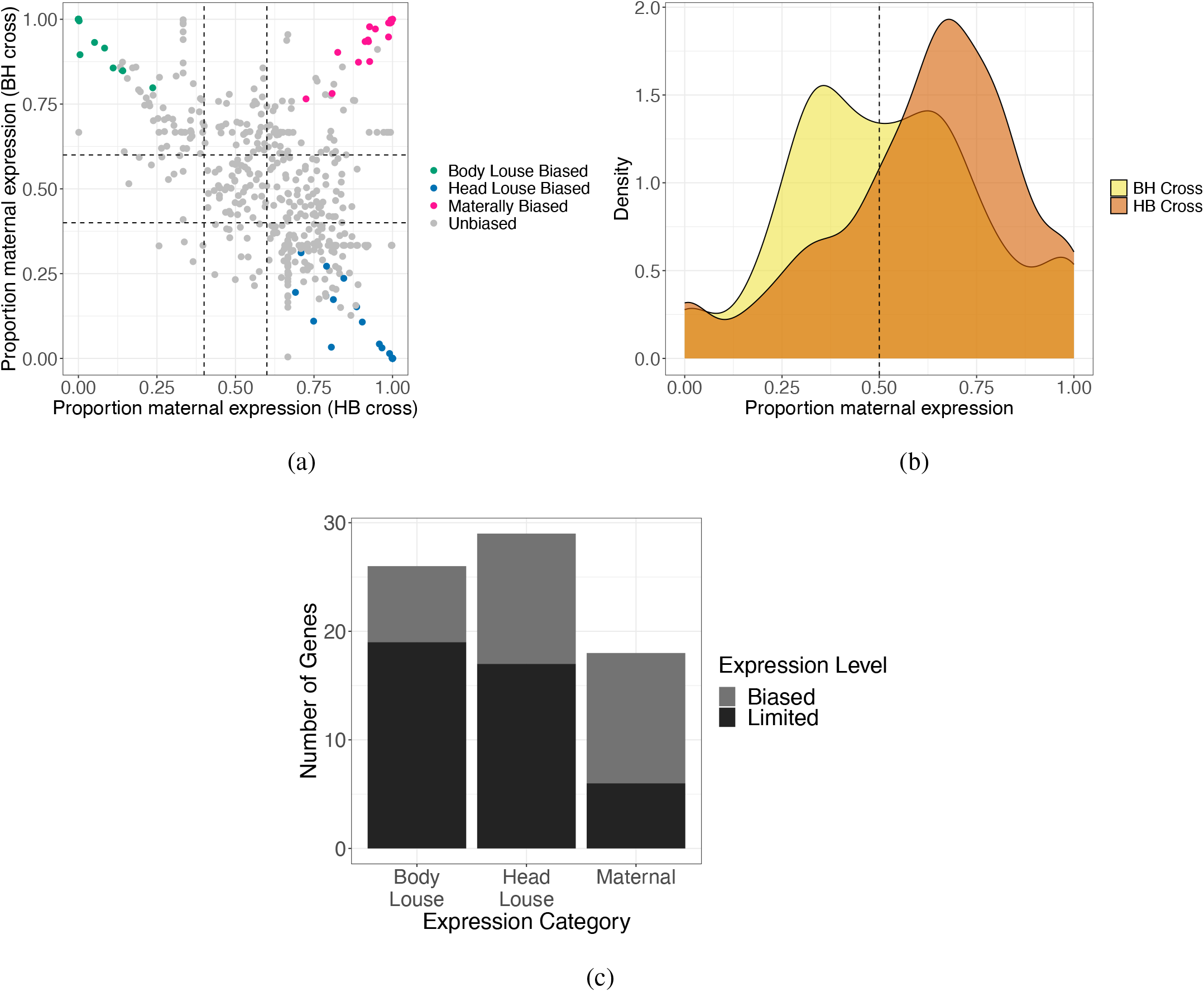
(a) Scatter plot showing the proportion of maternal expression for each cross direction for all genes tested (n=482). Each dot is a gene. The black lines represent 0.4 and 0.6 expression proportions, these were used as a hard cut-off in addition to a significant p-value in order to call a gene biased. (b) Density plot showing the distribution of genes by maternal expression proportion for each cross direction (n=482). BH = female body louse x male head louse, HB = female head louse x male body louse. (c) Staked bar plot showing the number of genes with expression bias in the reciprocal crosses. Biased genes are those with a significant maternal/paternal or head/body lice allelic ratio in both cross directions, corrected p-value <0.05 and biased expression proposition >0.6. Limited genes meet the above criteria however, they show complete silencing of one allele, i.e. expression is limited to either the maternal / head louse / body louse allele.

A GO enrichment analysis of the maternally biased and maternally limited genes reveals multiple functions, specifically many processes are associated with microtubule and actin filament organisation (supplementary 1.0.3), with the GO terms “*meiotic spindle organization*” (GO:0000212), “*establishment of mitotic spindle orientation*” (GO:0000132) and “*mitotic spindle elongation*” (GO:0000022) specifically enriched. The segregation of maternal and paternal chromosomes during spermatogenesis is a key feature of PGE and is expected to be under maternal control. To explore this further, we identified Drosophila orthologs for the maternally biased genes and we find the gene responsible for the enrichment of “*meiotic spindle organiszation*” (*xmap215*) is an ortholog of the *mini spindles* gene in Drosophila (FBgn0027948, supplementary 1.0.4). This gene is involved in microtubule organisation during both mitosis and meiosis (Brittle and Ohkura, 2005; Moon and Hazelrigg, 2004). The enriched GO terms, and the identification of *xmap215* as maternally biased, suggest a role for some of the maternally biased/limited genes in the segregation of the maternal and paternal chromosomes during spermatogenesis. However, it is worth noting that only around 43% of all annotated genes within the current *P. humanus* genome have associated GO terms, as such we are missing functional information for some of the maternally biased/limited genes.

### Ecotype-specific expression in F1 hybrid offspring

In additional to identifying parent-of-origin gene expression we were also able to identify ecotype-of-origin expression in the above 482 genes (supplementary 1.0.2). We find 26 genes show higher expression from the body louse alleles compared to the head louse alleles in the hybrid male offspring, 19 of which show complete silencing of the head louse allele (Figs. 2a and 2c). We see a similar number of genes showing head louse allele expression bias, 29 total, with 17 of these being completely silenced from the body louse allele (Figs. 2a and 2c). Interestingly, we also see a large skew generally for higher head louse allelic expression (Fig. 2b).

A GO enrichment analysis of these genes shows significant body louse biased/limited genes are involved in many cellular regulatory processes and metabolic processes, including “*regulation of meiotic cell cycle*” (GO:0051445) (supplementary 1.0.3). Whereas significant head louse biased/limited genes are more enriched for developmental processes, including “*adult somatic muscle development*” (GO:0007527), “*oligodendrocyte development*” (GO:0014003) and “*cellular component assembly involved in morphogenesis*” (GO:0010927) (supplementary 1.0.3). As with above, some of the biased/limited genes do not have GO annotations and so some functional information is missing.

Finally, we also checked to see if the genes with ecotype-specific bias in hybrid males are differentially expressed between pure head and body lice males. We find only one gene which shows both ecotype-specific expression in hybrid males and is also differentially expressed between pure ecotypes. *Programmed cell death protein* (PHUM549420) shows higher head lice allele expression in hybrid males but higher expression in pure body lice males compared to pure head lice males.

## Discussion

In this study we used RNA-Seq from male hybrid offspring of head and body lice ecotypes of *P. humanus* to identify both parent-of-origin and ecotype-of-origin gene expression patterns. We find that hybrid offspring show mostly biparental expression of genes, meaning that the paternal chromosomes of males are not transcriptionally silent in *P. humanus*, contrary to what is seen to some extent in other species with PGE. We find no genes with paternal expression bias, however we do find several genes with maternal expression bias, some of which are involved in meiosis and chromosome segregation, which means that they could play a role in the elimination of the paternal chromosomes during spermatogenesis. We also find several genes which are expressed in an ecotype fashion, i.e. either the head/body louse copy is always more expressed in hybrid offspring compared to the other copy. We find these ecotype-specific expressed genes in hybrid offspring are mostly not differentially expressed between adults of the pure ecotypes. Finally, we have identified a potential role for epigenetic processes in ecotype differences, as indicated by differentially expressed genes between ecotypes being enriched for epigenetic-related GO terms.

### Transcriptionally active paternal chromosomes in males

The identification of full biparental gene expression in *P. humanus* is the first documented case in a species with PGE. Recent work in the mealybug *Planococcus citri*, which also displays PGE, used RNA sequencing to show almost no expression of the paternal genes in males (de la Filia *et al*., 2021). In de la Filia *et al*. (2021), a few genes did show some level of paternal expression, meaning they escape full silencing, however this was less than 1%. In another scale insect clade (Ericoccidae) there is variability in the presence of paternal chromosome heterochromatinization and the degree of paternal gene expression even between closely related species (Hodson *et al*., 2022). This finding suggests that patterns of male gene expression and ploidy might show substantial turnover within scale insects. This situation might also be the case in lice where PGE is found across all parasitic lice and one genus of booklice (*Liposcelis*) that is likely the sister clade to the parasitic lice. While all parasitic lice studied so far lack paternal chromosome heterochromatinization. *Liposcelis* species do exhibit paternal chromosome heterochromatinization (Hodson *et al*., 2017). It is currently not clear what drives transitions between biparental and uniparental (maternal) expression in males under PGE. The current leading hypothesis is that there is ongoing intra-genomic conflict between the paternal and maternal genome over transmission in males. Paternally-inherited alleles might be selected to escape elimination during spermatogenesis, while maternally-inherited alleles could be selected to avoid this by transcriptional silencing of paternal alleles (Herrick and Seger, 1999; Ross *et al*., 2010). It is possible, however, that there are selective drivers such as sexual-antagonistic selection for ploidy, but these have yet to be explored in the context of PGE.

It is also worth noting that the biparental expression we have identified appears to be skewed towards higher expression of maternal alleles generally (with the exception of head lice biased expression). This is unlikely due to mapping bias to the reference genome as we employed strict criteria to avoid this. Additionally, the reference genome was created from a body lice sample. As such, any mapping bias would favour reads from body lice alleles. We speculate that this general maternal bias may be caused by paternal silencing in some tissues but not others. Another possibility is that the mechanisms to silence paternal alleles are not consistently successful across the paternal chromosomes. This could lead to reduced levels of paternal expression without complete silencing. The application of single-cell RNA sequencing to this system would help to determine the consistency of maternal allele expression bias across tissues.

In addition to mostly biparental gene expression, we have also identified a number of genes which show significant maternal expression bias or completely limited expression from the maternal copy. The presence of a fraction of imprinted genes in human lice was anticipated: in the absence of somatic adaptations to prevent or reduce expression of paternal alleles, maternally imprinted genes can be expected to be involved in reproductive functions under PGE, chiefly during spermatogenesis. Faithful segregation of paternal and maternal chromosomes in meiosis is key to the evolution and maintenance of PGE. Lice display a highly modified achiasmatic meiosis, followed by several rounds of mitotic division at the end of which an asymmetrical division produces functional spermatids carrying the maternal genome, and degrading nuclei carrying the paternal genome. Differential segregation most likely occurs during the last of the series of mitotic divisions preceding the formation of active spermatids (McMeniman and Barker, 2006; de la Filia *et al*., 2018). This stage is a clear candidate for maternally-controlled segregation and elimination of parental chromosomes. In this context, the identification of *xmap215* as maternally biased is particularly intriguing. The Drosophila ortholog of this gene (*mini spindles*) has been shown to be involved in both stabilising and destabilising microtubles (Brittle and Ohkura, 2005), which could be important for the differential segregation of the maternal and paternal chromosomes. Whilst usually the mini spindle protein localises at centromeres (Lee *et al*., 2001), it can bind to non-centromere regions (Deng *et al*., 2021). Given that *P. humanus* is holocentric (Bressa *et al*., 2015) this adds to the possibility for a role of *xmap215* in paternal genome elimination. Whilst this is purely speculative, the identification of *xmap215* as a possible candidate for the regulation of PGE during spermanogenesis enables future functional studies which can knock-down this gene to examine its role in PGE.

Another expected category of maternally imprinting genes under PGE are those involved in tagging and expression control of paternal alleles. In mealybug and sciarid flies, paternal chromosomes display differential patterns of epigenetic marks, such as histone modifications (Goday and Ruiz, 2002; Bongiorni and Prantera, 2003; Khosla *et al*., 2006; Escribá *et al*., 2011). Whilst the GO enrichment analysis of the maternally biased genes identified here does not show any functions related to epigenetic processes, it is worth noting we were only able to assign GO terms to around <50% of these genes. Therefore, many functions of these genes remain unknown. Nevertheless, *P. humanus* does have a functioning DNA methylation system. DNA methylation appears to be involved in the silencing of paternal chromosomes in male mealybugs (although it is not clear if this is a cause of consequence of PGE) (Bain *et al*., 2021). An improved genome annotation for *P. humanus*, in addition to the identification of epigenetic differences between the maternal and paternal chromosomes would aid in the identification of the role of maternally-mediated epigenetic labelling in this system.

These findings provide the required groundwork for the identification of specific genes which may function to direct PGE within *P. humanus*. Future work utilising low input methods, such as single cell RNA sequencing, would allow screening for maternally expressed genes specifically in male gonad tissue where PGE takes place.

### Male ploidy and the evolution of resistance

The rate at which species respond to natural selection determines how fast they are able to adapt to changing environments. This process is relevant in lice in order to predict the evolution of resistance to newly introduced pediculicides used to treat louse infestations. The rate of adaptation is likely dependent on a species’ reproductive genetics. There has been some suggestion previously that the hemizygous expression in males under haplodiploidy and PGE could contribute to an increased rate of adaptation as rare recessive beneficial mutations are exposed to selection (Klein *et al*., 2021). However, we show that this is unlikely to occur in lice as males are fully diploid. Instead response to selection in males is likely impeded as males express a diploid set of genes but only pass on a haploid complement. Recent theoretical work shows that this leads to a reduction of the invasion probability of new alleles, particularly for male-function genes (Klein *et al*., 2021). This suggests that any control strategy aimed at males specifically might be more “evolution proof”, by reducing the speed with which resistance can evolve.

### Ecotype-specific gene expression

In addition to maternally biased genes we were also able to identify genes which show ecotype-specific expression. We find similar numbers of genes showing significant head louse expression bias to those showing body louse specific bias, with a large proportion of both having limited (i.e. monoallelic) expression. We also find higher expression generally from the head louse allele across genes. These asymmetric expression patterns are probably caused by cis-acting regulatory variants favoring the expression of one allele in both reciprocal crosses (Pollard *et al*., 2008; Wang and Clark, 2014). A previous study using reciprocal crosses of two related wasp species also identified a large number of species-specific expression bias in hybrid offspring (Wang *et al*., 2016). However, reciprocal crosses within species of honeybee (Galbraith *et al*., 2016) and bumblebee (Marshall *et al*., 2020), revealed little to no line-specific expression. The specific status of head and body lice has been long debated, but mounting evidence supports that their phenotypic differences are due to ecological factors and even their subspecific status has been questioned (Light *et al*., 2008). Currently, they are considered different ecotypes of *P. humanus*: body lice are believed to emerge regularly from head louse populations by colonising new breeding grounds in human clothes (Li *et al*., 2010). However, the identification of genes with ecotype-specific expression in hybrids is suggestive of a divergence between the two ecotypes.

We used genetic differences (i.e., SNPs) between ecotypes to identify parent/ecotype-specific gene expression. Previous work has identified very few genetic differences between head and body lice (Li *et al*., 2010). The low levels of divergence between the ecotypes meant that we were only able to assess parent/ecotype-specific expression in 482/10,992 annotated genes due to the low occurrence of differentiating SNPs between ecotypes within gene bodies. Whilst there appear to be very few genetic differences between ecotypes, differences in alternatively spliced transcripts have been observed (Tovar-Corona *et al*., 2015). We did not have the resolution to examine differences in alternative splicing in our study, however, future work to examine the possibility of parent/ecotype-specific alternative splicing could be fruitful in this system. We did, however, find a few differentially expressed genes between ecotypes. Previous evaluations of differentially expressed genes between pools of head and body lice from all developmental stages (instead of adult males only) are available, ranging from as low as 14 genes (Olds *et al*., 2012) to 552 genes (Previte *et al*., 2014). One of the most differentially-expressed genes in our study, which shows strong upregulation in head compared to body lice (or downregulation in body lice compared to head lice) is the immunity gene *Defensin 1*, part of the Toll pathway. This is consistent with previous studies (Kim *et al*., 2012) and is of interest given the different ability of the two ecotypes to vector the bacterial pathogen *Bartonella quintana*. We also found that differentially expressed genes are enriched for epigenetic related processes. DNA methylation has previously been implicated in alternative splicing in the honeybee (Flores *et al*., 2012). Epigenetic differences between ecotypes could explain the differences in life history and also the lack of need for genetic differentiation. Additionally, given that PGE is epigenetically mediated in other systems, future work focusing on epigenetic differences between head and body lice and maternal/paternal chromosomes in males is needed to better understand these systems.

## Conclusion

We have shown that adult males of *P. humanus* display biparental gene expression, which constitutes the first known case of a species with PGE in which genetic activity of paternal chromosomes in the soma is not affected by embryonic heterochromatinization or (partial or complete) elimination. The absence of adaptations to silence the paternal genome as a whole in the human louse can facilitate unmasking genes involved in PGE (either enforcing it when maternally imprinted or combating elimination when paternally imprinted), most specifically during spermatogenesis. For example, the identification of the maternally-biased *xmap215* gene lends itself to future functional studies of the mechanisms of PGE. Finally, the identification of ecotype-specific expression in hybrids of head and body lice, in the context of low genetic diversity, is suggestive for a role of epigenetic processes in ecotype differences.

## Acknowledgements

This work was funded by a European Research Council Starting Grant: PGErepro, NERC fellowship NE/K009516/1, and Dorothy Hodgkins fellowship DHF\R1\180120 awarded to L.R. A.F. was supported by the Darwin Trust of Edinburgh. H.M. was supported by a Leverhulme Trust Research Project Grant: RPG-2020-363, awarded to E.B.M. This research used the ALICE2 High Performance Computing Facility at the University of Leicester.

## Author contributions

A.F., L.R. and J.M.C. conceived the study. A.F. and R.C. conducted the experiment. A.F., H.M. and E.B.M. analysed the data. A.F. and H.M. wrote the initial manuscript. All authors contributed to and reviewed the final manuscript.

## Data Accessibility

Data has been deposited in GenBank under NCBI BioProject: PRJNA968062. All code is available at: https://github.com/MooHoll/head_and_body_lice_imp.

