## supplementary 2 for "Lack of paternal silencing and ecotype-specific expression in head and body lice hybrids"

### Supplementary 2.0: additional figures

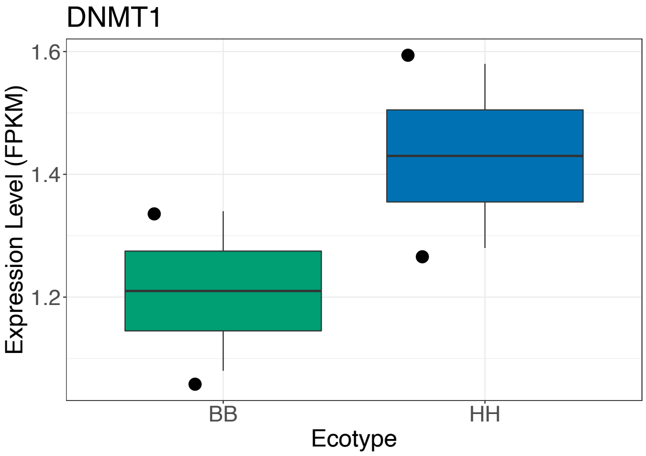


(a)

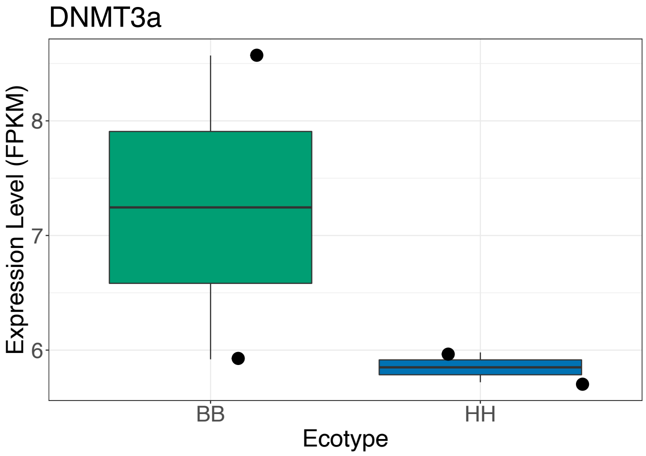


(b)

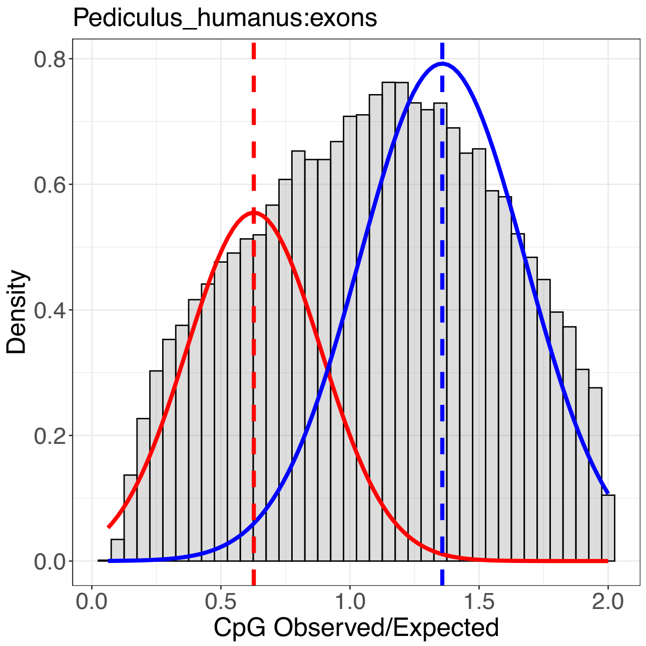


(c)

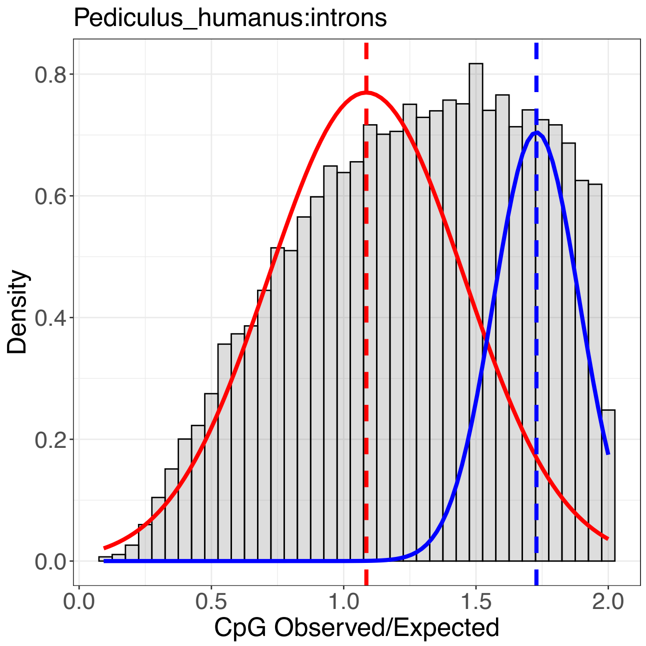


(d)

Figure S1: (a and b) Boxplots of expression levels of two DNMT genes. BB refers to body lice and HH refers to head lice. (c) CpG observed/expected distribution for all exons in the *P. humanus* genome and for introns (d). CpG o/e is the ratio of the observed number of CpG sites in a given region compared to the expected number of CpG sites, given the GC content of that region. A depletion of CpG sites (CpG o/e $< 1$) is indicative of DNA methylation presence as DNA methylation degrades CpG sites over time due to the increased deamination of cytosines to thymines. Species with DNA methylation tend to show two CpG o/e peaks, one around 0.5 and another around 1.0, representing methylated and unmethylated regions.
